# An effective primary and memory CD8^+^ T cell response is directed towards an H-2K^b^ restricted HSV1-gB-SSIEFARL epitope in the experimentally infected zebrafish

**DOI:** 10.1101/2025.08.14.670278

**Authors:** Dhaneshwar Kumar, Syed Azeez Tehseen, Sharvan Sehrawat

## Abstract

We sought to identify and characterize HSV1 specific CD8^+^ T cells in the experimentally infected zebrafish (zf), a model organism that offers real time tracking of cellular dynamics. We generated a fluorescently labelled class I MHC tetramer of zebrafish using an H-2K^b^ restricted peptide of gB protein of HSV1 (Uda-HSV1-gB-SSIEFARL-tetramer) for analysing the differentiation of virus-specific CD8^+^ T cell response. We show that the Uda-SSIEFARL-tetramer^+^CD8^+^ T cells rapidly expand in the lymphoid organs and are efficiently recruited to non-lymphoid tissues of HSV1 infected zebrafishes. The expanded cells upregulate effector cytokines; IFN-γ, IL-2, TNF-α and cytolyze SSIEFARL-peptide pulsed targets. Uda-SSIEFARL-tetramer^+^CD8^+^ T cells efficiently migrate to the infected tissues as demonstrated by fluorescent microscopy and *in vivo* imaging. The Uda-SSIEFARL-tetramer^+^ memory cells are recalled in the response following a secondary infection of zebrafish with HSV1. We, therefore, make a rather intriguing observation that zebrafishes infected with HSV1 generate an efficient CTL response against H-2K^b^ restricted HSV1-gB-SSIEFARL epitope despite their evolutionary divergence from mice ∼445 million years ago. We also argue that zebrafish could serve as a better accessible model organism for studying *in vivo* dynamics of virus-specific CD8^+^ T cells.

## Introduction

The anatomical visual transparency of zebrafish offers analysis of immune cell dynamics in ways not possible in other vertebrates and therefore this model organism could unearth novel aspects of pathogenesis. Kidney marrow and pancreas serve as the major sites of haematopoiesis and most of the haematopoietic cells have been demonstrated to exist in zebrafish using genetically tractable fluorescent markers^1–4^. Immune cells expressing Foxp3, a master transcription factor which confers to the cells suppressive functions, were demonstrated but the studies aimed at investigating the differentiation of helper and cytotoxic T cells have not been performed in zebrafish.

Zebrafish as a model organism has provided key insights into the developmental and regeneration biology, toxicology, haematopoiesis as well as immunoinflammatory reactions caused by some microbes^5–14^. The functionally and morphologically mature adaptive immune system is evident at 4-6 weeks after fertilization and hence the larvae of zebrafish survive with only the innate immune responses^15,16^. This temporal distinction of innate and adaptive immune parameters can help better understand the relative contribution of two branches of immunity during pathophysiological responses without resorting to depletion studies. Dendritic cells, γ8 T cells and Treg-like cells were demonstrated to exist in zebrafish^17–23^. However, the studies aimed at investigating anti-viral adaptive immune response are largely lacking. This is predominantly attributed to the unavailability of reagents to detect antigen-specific T cells. Backed by bioinformatic analysis, we generated a class I MHC (Uda) tetramer loaded with an immunogenic epitope of HSV1 in C57BL/6 mice and used it in an attempt to measure the response of antigen-specific CD8^+^ T cells in the experimentally infected zebrafish as a primer investigation. We observe an efficient expansion of tetramer positive cells followed by their contraction which left behind a fraction of the expanded cells formed memory. The recallability of memory cells was demonstrated in the HSV1 infected zebrafishes. The expanded cells efficiently cytolyzed the peptide-pulsed targets and upregulated the effector cytokines. The observations underscore the existence of conserved antigen processing and presentation mechanisms in zebrafish and mice despite their evolutionary divergence ∼445 million years ago. This study also paves the way for optimally utilizing zebrafish as a model for elucidating not only the differentiation pathways of antigen-specific CD8^+^ T cells but also their role in viral pathogenesis.

## Results and Discussion

### 1. Experimentally infected zebrafishes mount HSV1-specific immune response

The animals were inoculated with 2×10^6^ pfu of HSV1 via intraperitoneal route and the virus loads were measured in the homogenates of spleen and the rest of their body parts collected at different time post infection from the euthanised animals (Fig 1A). The infectious viral particles were recovered from the splenic tissues between 24 and 48 hrs post infection (hpi) (Fig 1B and C). The virus was also recovered from the homogenates of body parts of zebrafishes that were devoid of splenic tissues for upto 72hpi (Fig 1B and C). We also measured the mRNA expression of one of the essential genes of HSV1, the catalytic unit of DNA polymerase (UL30), in the infected zebrafishes to ascertain the *in vivo* replication of virus following inoculation. The expression levels of UL30 increased by 5-fold at 24hpi and reached to a basal level by 72hpi (Fig D). Herpesviruses are known to adopt two modes of life style viz. lytic cycle and latency^24,27^. Following a primary infection, HSV1 replicates in the host and resort to a latency in neuronal tissues once the replication phase is over ^26,27^. Many aspects of the establishment, maintenance and reactivation of HSV1 from latency remain to be fully investigated for the lack of a reliable and relevant animal model. We, therefore, investigated whether or not HSV1 infection establishes latency in zebrafishes by measuring the expression of a subunit of DNA polymerase (UL30) and latency associated transcripts (LATs) in spleen and brain tissues. The expression of UL30 and LATs could indicate the replication and latency, respectively. We observed ∼3 and 5-fold increase in the expression of UL30 in spleen and brain tissues of HSV1 infected animals at 24hpi (Fig 1E). Thereafter, the expression was reduced and by 72hpi, it reached at basal levels in both the tissues (Fig 1E). The expression of LATs was increased by 80-fold at 24hpi in brain but not in the spleen splenic tissues. Thereafter, their expression decreased but still remain evident by 3dpi in the brain tissues (Fig 1F). That the LATs were expressed in the neuronal tissues of HSV1 infected zebrafishes could pave the way for assessing host-virus interaction in HSV1 infected zebrafishes and measuring the contribution of virus-specific CD8^+^ T cells in the viral control. In some of the earlier studies HSV1 infection was performed in zebrafish^28–30^. While our results support the published results, we additionally demonstrate the presence of replicating virus particles in the spleens and elsewhere in the infected zebrafishes^21^. The susceptibility of zebrafish larvae for ocular HSV1 infection was enhanced by liposome encapsulation of the virus HSV1^31^.

**Figure 1.**
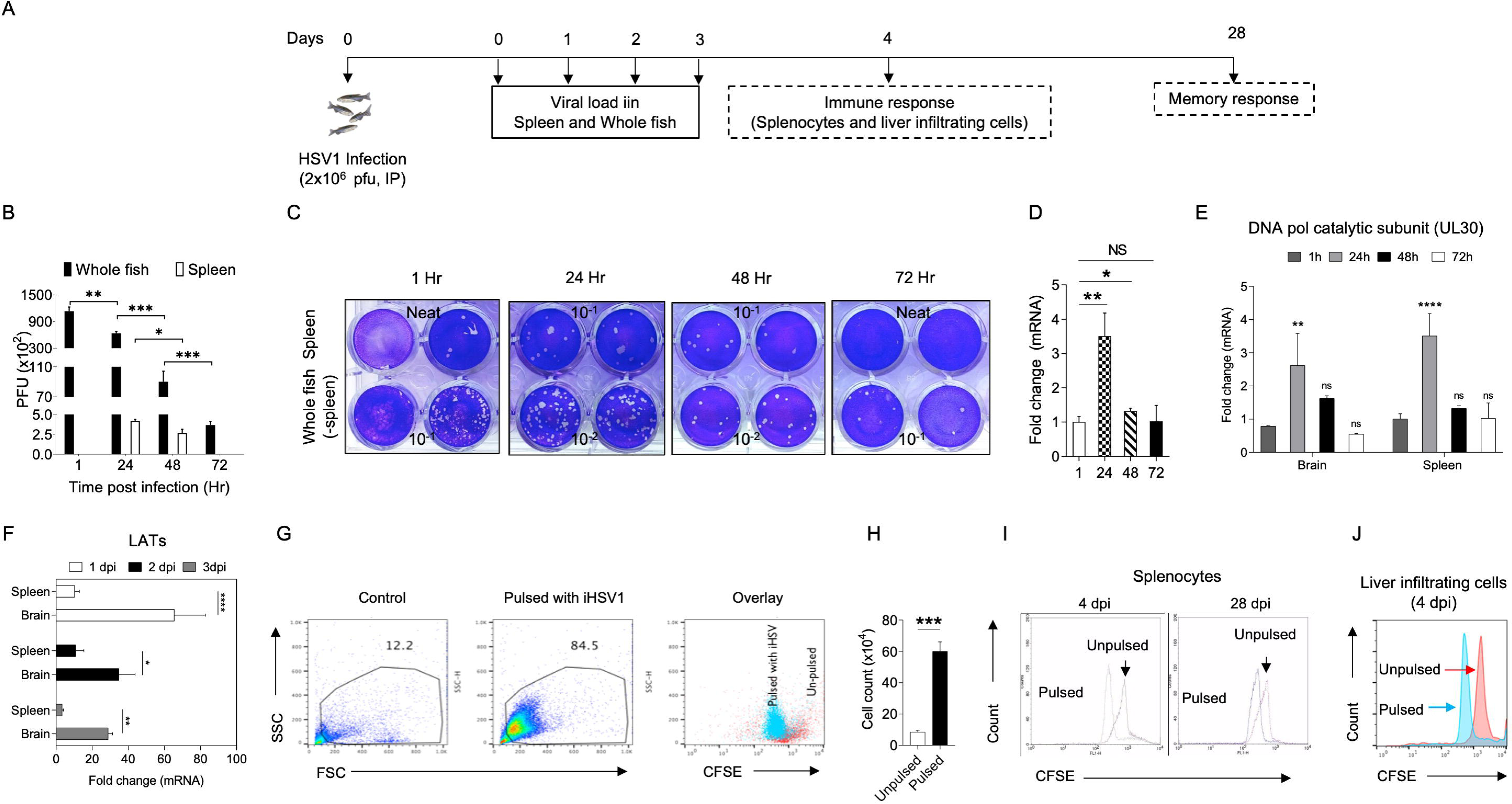
Experimentally infected adult zebrafishes permit HSV1 replication and induction of specific immune reactivity. A. In-house bred zebrafishes were i.p. infected with 2×10^6^ pfu of HSV1-KOS. Spleen and the remaining body parts were collected from euthanised animals at 0, 1, 2 and 3dpi. The tissue homogenates were prepared and the supernatants were used for quantifying the viral load by plaque forming assays. B. Bar diagrams show the viral loads in spleen samples and the remaining body parts of zebrafishes collected at indicated time post HSV1 infection. C. Representative images of developed plates used for quantifying the viral loads by plaque forming assays are shown. D. RNA was isolated from splenic tissues of HSV1 infected zebrafish at different time post infection and converted into cDNA. The expression levels of catalytic unit of DNA polymerase (UL30) of HSV1 is shown as fold change. E. Bar plots show the mRNA expression of catalytic subunit of HSV1 DNA polymerase (UL-30) by qRT-PCR from brain and spleen samples at indicated time are shown. F. The expression levels of HSV1 latency associate transcripts (LATs) in spleen and brain tissues of HSV1 infected zebrafish is shown by bar plots. The experiments were repeated three times using four zebrafishes at each time and the results from representative experiments are shown. Results were analysed by one way ANOVA. The level of significance are represented as *, p<0.05, **, p<0.01 and ***<0.001. G. HSV1 infected zebrafishes were sacrificed at 4 or 28dpi to isolate spleen and liver tissues. The prepared single cell suspensions were CFSE labelled and stimulated with the UV-inactivated HSV1 at 1MOI for assessing the virus induced proliferative potential of splenocytes. Representative FACS plots show the relative abundance of lymphocytes in unstimulated and the stimulated cells. Overlaid FACS plots show the levels of CFSE dilution in two groups of animals after 72 hrs of culture. H. Bar diagram show the number of cells in control and stimulated samples. I. Representative overlaid histograms show the proliferation of virus antigen stimulated zebrafish splenocytes in the acute phase at 4dpi and memory phase at 28dpi. J. The proliferation of liver infiltrating lymphocytes stimulated with viral antigens and controls is shown at 4dpi is shown by overlaid histograms. Student ‘t’ test was used to determine the level of statistical significance between different groups. Mean ± SEM values are shown. The level of significance are represented as *, p<0.05, **, p<0.01 and ***<0.001.

To assess the responsiveness of immunocytes to HSV1 infection, we CFSE labelled splenocytes from the infected zebrafishes and the labelled cells were stimulated from individual zebrafish in separate wells with the inactivated HSV1 (iHSV1). Unstimulated cells served as the control. The cells were analysed for their scattering properties, relative abundance and the dilution of CFSE 72hrs later (Fig 1G). We observe an approximately seven-fold increase in frequencies of the expanded immune cells in the HSV1 pulsed samples as compared to the no virus control (84% in iHSV1 pulsed splenocytes vs ∼12% in control) (Fig 1G and H). Furthermore, a large majority of HSV1 pulsed cells diluted CFSE content which suggested for their extensive proliferative response (Fig 1G and H). We also measured the proliferation of splenocytes and liver infiltrating lymphocytes obtained from HSV1 infected zebrafishes in the acute (4dpi) and memory (28dpi) phase (Fig 1I and J). The CFSE labelled liver infiltrating immune cells as well splenocytes obtained from HSV1 infected zebrafishes divided extensively in response to restimulation with HSV1 antigens when analysed *ex vivo* in the acute (4dpi) and memory (28dpi) stage of infection (Fig 1I and J, data not shown).

We show that the infected zebrafishes allow HSV1 replication and expand immunocytes in both the splenic and liver tissues. The immune cells recovered from HSV1 infected animals extensively proliferate *ex vivo* upon restimulation with the viral antigens. Furthermore, the HSV1 adopts to a latent life cycle in the neuronal tissues of the infected zebrafish.

### 2. Generation of fluorescent MHC class I (Uda) tetramer of zebrafish for detecting HSV1 specific CD8^+^ T cell

We first aligned the sequences of heavy chain of U and Z lineages of class I MHC molecule of zebrafish with that of a mouse H-2K^b^, the later displays one of the major immunodominant viral peptide of gB_498-505_ (SSIEFARL) of HSV1^32,33^ (Fig 2A). U lineage class I MHC molecules of zebrafish and H-2K^b^ formed one cluster while the Z lineage formed a separate cluster (Fig 2A). We, therefore, aligned the top five U lineage sequences of heavy chain i.e., Ula, Uma, Uda, Uaa and Uba with those of H-2K^b^ to assess their homology and identity in the α1 and α2 domains^34^. H-2K^b^ showed 36% and 41% sequence identity and 55% and 58% structural homology at an RMSD value of 1.553Å with Uaa and Uda of zebrafish, respectively (Fig 2A-E). We then searched for potential contacts in the peptide binding pockets of Uda heavy chains that might interact with the anchors of SSIEFARL peptide (Fig 2B and C). Residues V76, T143, K146 and W147 of H-2K^b^ and V74, T140, K143, W144 of Uda showed strong interaction with the anchor residue (L8) of SSIEFARL peptide. Similarly, residues V76, D77 and W147 of K-2K^b^ and V74, D75 and W144 of Uda interacted with R7 of the peptide (Fig 2B and C). The residues Y7, Y159, W167 and Y171 of H-2K^b^ and Y7, Y156, W164 and Y168 of Uda showed interaction with S1 and S2 of the peptide (Fig 2B and E). The docking of SSIEFARL peptide on an overlaid H-2K^b^ and Uda heavy chains revealed the presence of conserved interacting residues with the anchors of SSIEFARL peptide (Fig 2B-E). The results of our sequence analysis and the docking of SSIEFARL peptide on the overlaid H-2K^b^ and Uda revealed the presence of either the same amino acid or those with similar biochemical characteristics at the designated position in H-2K^b^ and Uda molecules (Fig 2B). These results suggested that Uda molecule could present gB-SSIEFARL peptide of HSV1. We, therefore, planned to generate a fluorescently labelled Uda tetramer refolded with the SSIEFARL peptide of HSV1 gB protein for detecting the virus reactive CD8^+^ T cells. We amplified β2-microglobulin and the extracellular domain of Uda with a biotin acceptor peptide (BAP) sequence at the C-terminus using the primers (Table 1 and Fig 3A and B). The BAP tag helps biotinylate the expressed Uda heavy chain in a site specific manner at lysine residue^35^. Refolding of the peptide, β2-microglobulin and the heavy chain of Uda can then generate a class I MHC monomer for zebrafish which can be tetramerised using a fluorochrome conjugated streptavidin for increasing its avidity of interaction with the T cell receptors to detect antigen-specific T cells. We cloned, expressed and purified Uda and β2-microglobulin from inclusion bodies (Fig 3B-H). We then refolded these proteins (Uda and β2-microglobulin) in the presence of SSIEFARL peptide by a rapid dilution method^36,37^. The refolded samples, fractionated using S200 column by size exclusion chromatography, yielded four peaks which were resolved using a 12% SDS-PAGE. Both the heavy chain of Uda (∼35kDa) and β2 microglobulin (∼13kDa) were present in the peak 2 which indicated the generation of the monomer (Fig 3I and J). The peptide being smaller in size could not be detected in the gel. We also observed refolding of the Uda heavy chain and β2 microglobulin with SIINFEKL, a peptide of Ovalbumin and FAPG(Anp)YPAL, a photocleavable cleavable peptides, both restricted by H-2K^b^ but the area under peak 2 was lesser as compared to that obtained with the SSIEFARL peptide (data not shown). To ensure the refolding and generation of a monomer, we analysed the pooled and concentrated fractions from peak 2 by circular dichroism and observed 47.11% of β-sheets and 3.66% of α-helical structures (Fig 3K). We then biotinylated the refolded monomers using BirA ligase, purified the biotinylated-monomer by gel filtration and detected using streptavidin-HRP (Fig 3L). We observed a band of ∼37kDa with the resolved monomer (Fig 3L).

**Figure 2.**
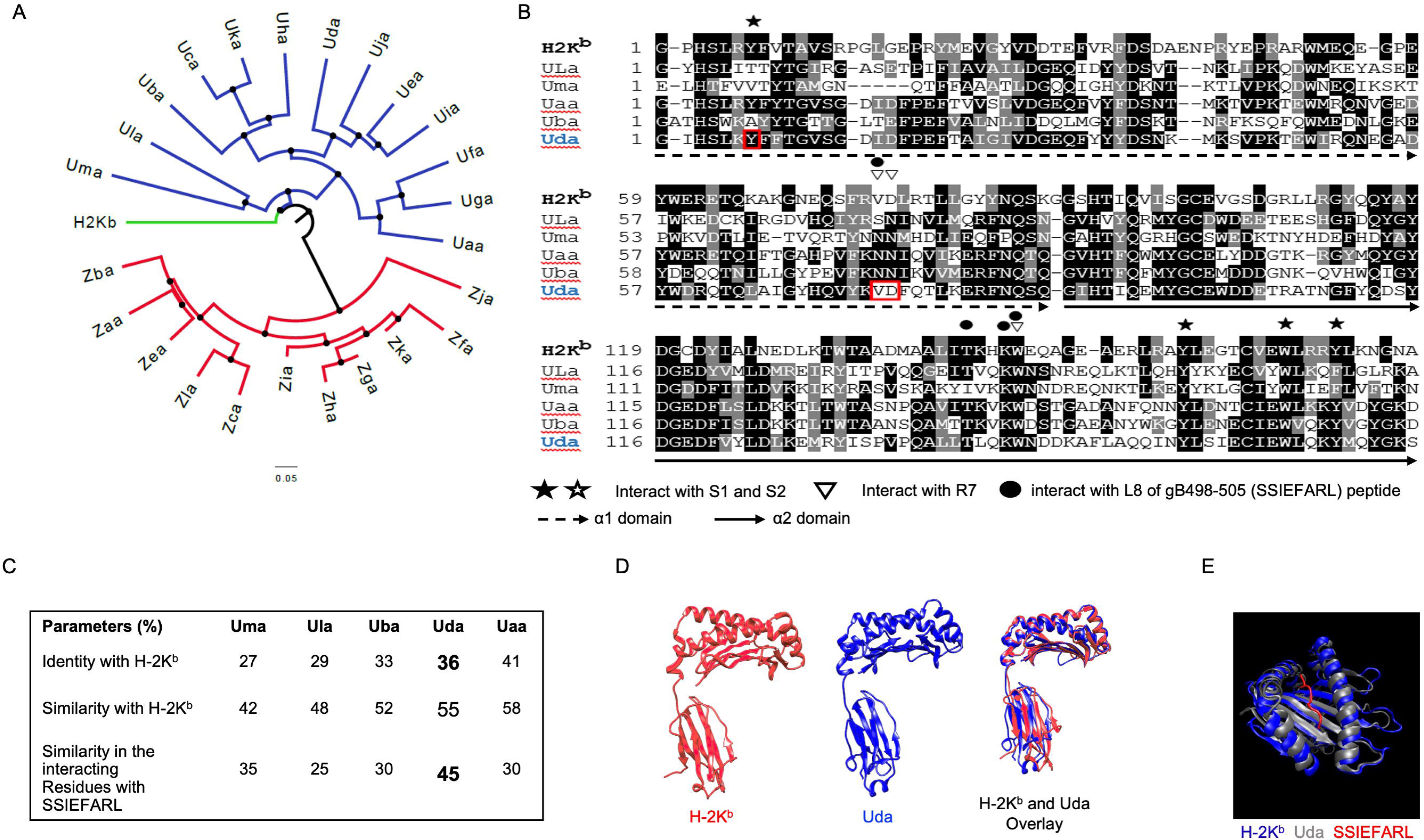
Identification of putative anchors for SSIEFARL peptide with class I MHC molecule (Uda) of zebrafish. A sequence alignment of different class I MHC heavy chains of zebrafish and H-2K^b^ was performed to search for homology and the anchor residues for SSIEFARL peptide. A. The generated phylogenetic tree demonstrates similarity between H-2K^b^ and heavy chains of zebrafish class I MHC molecule. B. Top five sequences of U lineage showing high homology with H-2K^b^ were analysed for identifying the amino acids which make contact with the anchors of HSV1-gB-SSIEFARL peptide. The shared amino acids of H-K^b^ and Uda molecules which showed interactions with SSIEFARL peptide are shown in bold red boxes. The α1 and α2 domains of Uda are shown by dotted and complete lines, respectively. C. Analysis of heavy chains of top five zebrafish U lineage which showed high homology with those of H-2K^b^ shows the percent identity and similarity. D. Putative structure of Uda compared to the crystal structure of H-2K^b^ heavy chain and their overlay show similar 3D structures as well as the peptide binding pockets for the cognate peptide. E. An overlaid image of putative Uda tertiary structure with H-2K^b^-SSIEFARL monomer is shown.

**Figure 3.**
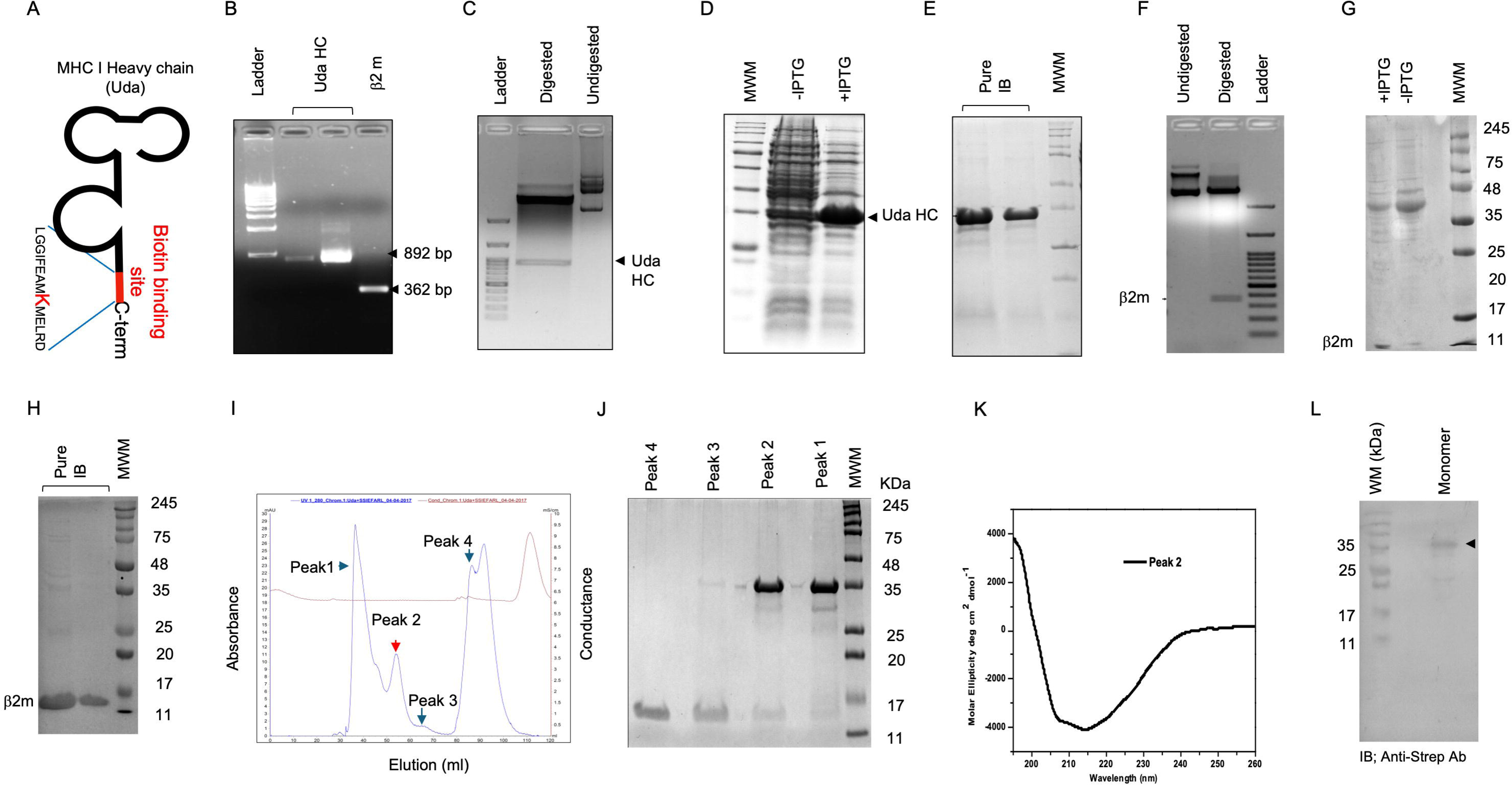
Generation of class I MHC tetramer for zebrafish for measuring and quantifying HSV1 specific T cells. For generating class I MHC tetramers for zebrafish, heavy chain of Uda and β_2_ microglobulin were cloned, expressed and purified. The purified proteins were refolded in presence of HSV1-gB-SSIEFARL peptide and the refolded monomers were purified using size exclusion chromatography. After biotinylation the monomers were fractionated and the purified monomers were tetramerized using streptavidin-PE. A. A scheme shows the modifications for producing the recombinant Uda. B. PCR amplification of Uda heavy chain and β_2_ microglobulin resolved using a 1% agarose gel is shown. The Uda heavy chain (C-E) and β_2_ microglobulin (F-H) were cloned in pET22b vector (C and F), expressed (D and G) and purified from the inclusion bodies (E and H). The expressed heavy chain and β_2_m proteins were purified from inclusion bodies and the purity was assessed by 12% SDS-PAGE (E and H). The purified proteins were refolded in the presence of SSIEFARL peptide and the monomers were purified by size exclusion chromatography using S200 Sephadex column. I. The chromatogram shows the elution profile of different polypeptides collected form the refolding buffer. J. The polypeptides present in different peaks were analysed by 12% SDS-PAGE. K. Circular dichroism of peak 2 protein (Uda monomers) was performed and the different peaks show the refolded structure. L. Western blotting was performed using streptavidin-HRP for measuring the biotinylation efficiency of purified monomer before tetramerization.

**Table 1.**
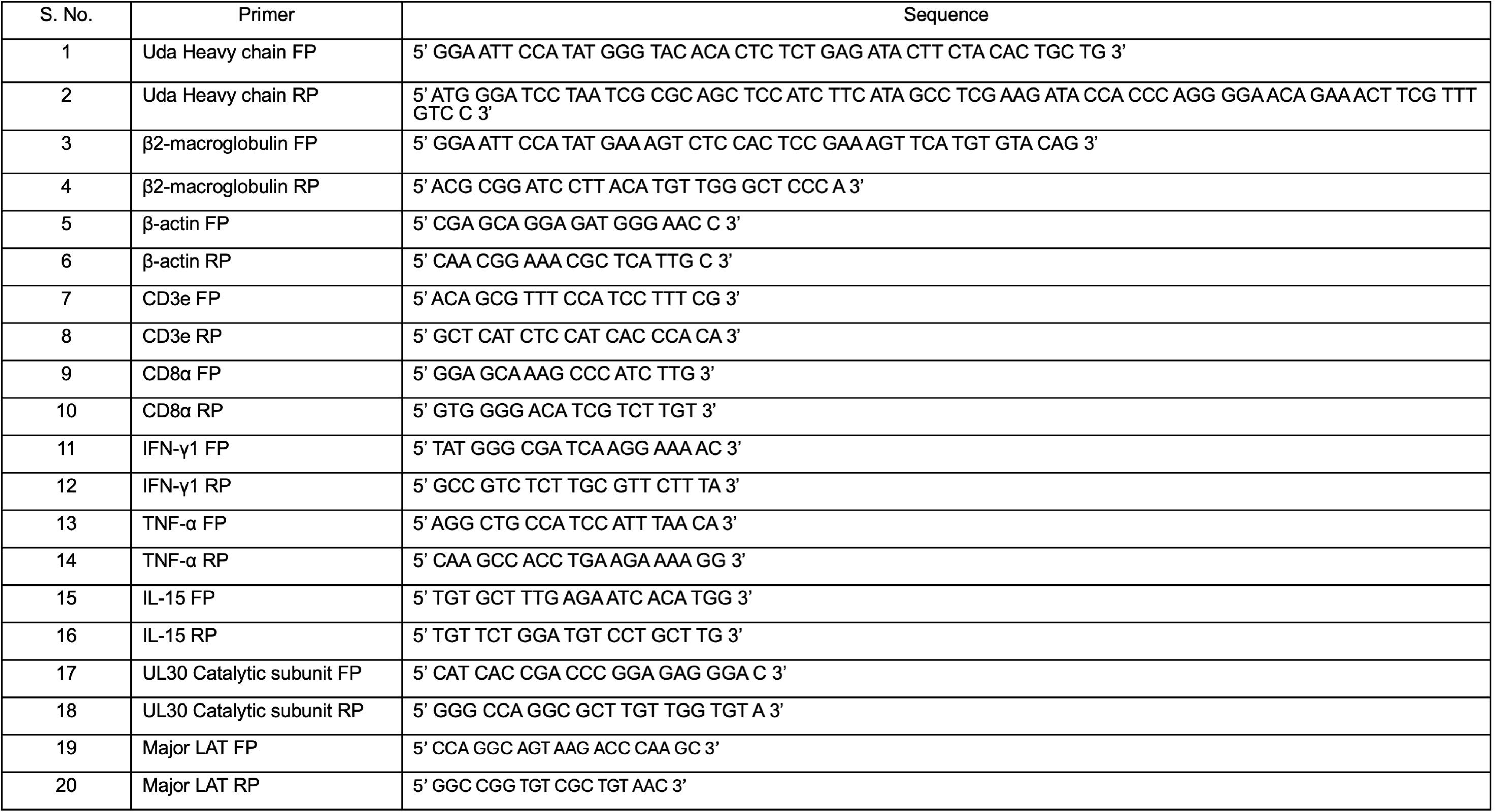
A list of primers used in the study is shown.

### 3. Measuring virus-specific CD8^+^ T cells during HSV1 infection of zebrafish

Uda-SSIEFARL-tetramers were used to detect and measure the kinetics of HSV1 specific CD8^+^ T cell response in the infected zebrafishes at 5dpi (Fig 4A). We detected ∼9% and ∼ 2% tetramer^+ve^ cells in HSV1 infected and the uninfected controls while an irrelevant peptide loaded tetramer (Uda-SIINFEKL-tetramer) showed a basal level (∼1%) of staining in the infected zebrafishes (Fig 4B, data not shown). A back gating of tetramer^+ve^ cells showed that ∼98% of such cells originated from the lymphocyte population based on the scattering properties (Fig 4B, right plot). These results confirmed the functionality of Uda-SSIEFARL-tetramers and demonstrated their ability to identify HSV1 reactive CD8^+^ T cells in the infected zebrafishes. We then analysed the kinetics of the expanded virus-specific CD8^+^ T cells in HSV1 infected zebrafishes and observed increased frequencies of tetramer^+ve^ cells until 5-6dpi (∼7% tetramer^+ve^ of total live cells) (Fig 4C and D). Thereafter, the frequencies of tetramer^+ve^ cells decreased and stabilized at ∼2.5% of total live cells by 45dpi (Fig 4C and D). Once a primary infection is controlled, the expanded CD8^+^ T cells undergo a contraction phase leaving behind a pool of memory CD8^+^ T cells that can be recruited in the response following a subsequent homologous infection. We, therefore, tested the recallability of the persisting tetramer^+ve^ cells following a secondary infection with HSV1. More than two fold increase in HSV1 reactive tetramer^+ve^ cells was recorded in the reinfected animals (Fig 4C and D). A relatively subdued recall response is routinely observed in animals re-infected with HSV1 and it could be due to the reduced levels of viral antigens resulting from the induced neutralizing antibodies. We also measured the *in vitro* proliferative response of the HSV1 expanded CD8^+^ T cells. The splenocytes were prepared from the HSV1 infected animals at 5dpi. The cells from individual zebrafishes were isolated and labelled with CFSE. The labelled cells were then cultured in separate wells in absence or the presence of SSIEFARL-peptide and the extent of proliferation was measured at the indicated timepoints (Fig 4E-F). The cells cultured without the peptide did not proliferate while those added with the peptide proliferated extensively at 72 and 96 hrs post incubation (Fig 4E-F). These results showed an antigen-specific expansion of immune cells.

**Figure 4.**
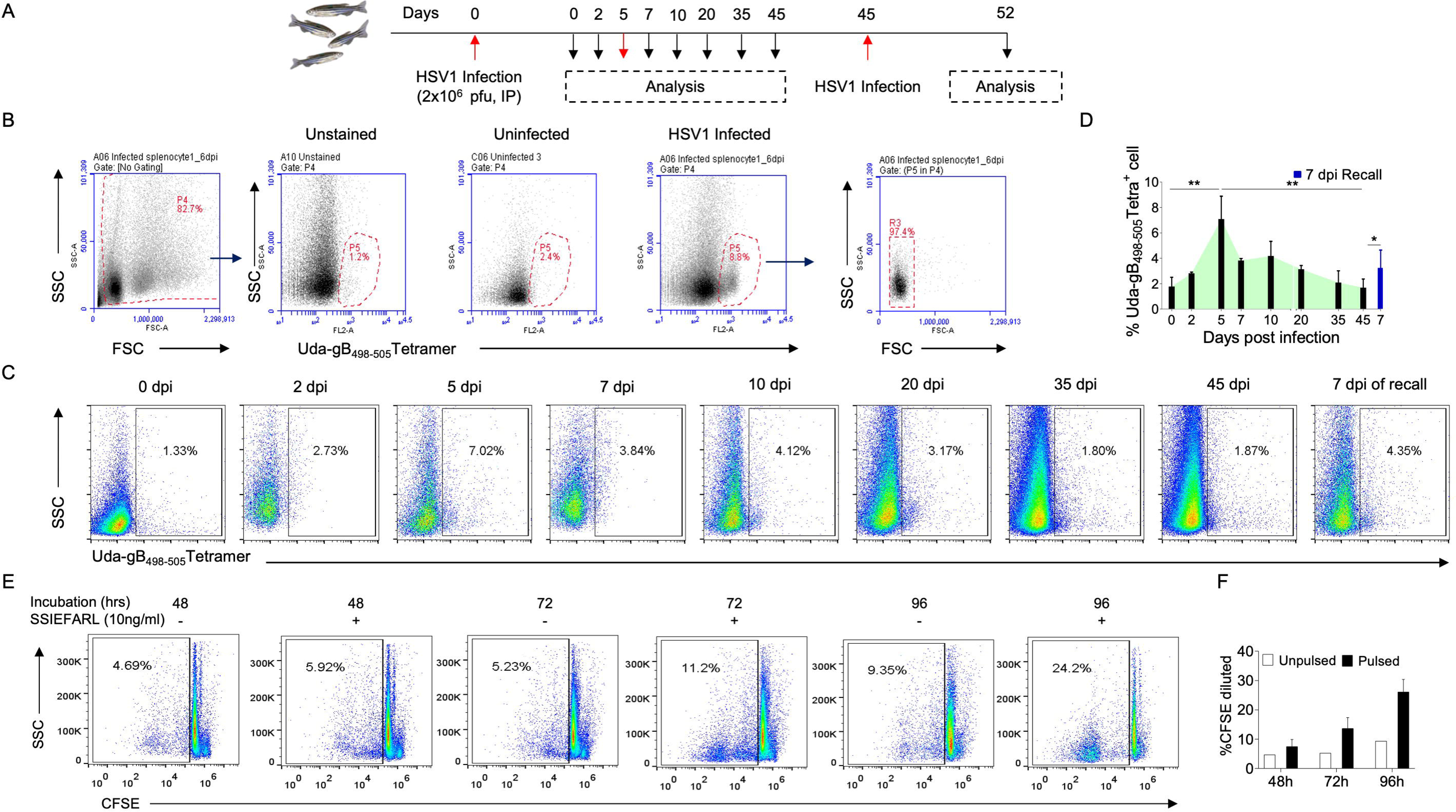
Detection of virus-specific CD8^+^ T cell response and measuring their kinetics in experimentally HSV1 infected zebrafish. A. Experimental plan for the detection and kinetics measurement of HSV1 specific CD8^+^ T cells in HSV1 infected zebrafishes (2×10^6^ pfu/zebrafish i.p.) is shown. Splenocytes were collected at indicated time points and stained with Uda-SSIEFARL-tetramers for detection of HSV1 specific T cells. B. FACS plots show the gating strategy of Uda-SSIEFARL-tetramer staining of splenocytes. Tetramer^+ve^ cells were then backgated to identify their origin and the scattering characteristics. C. Representative FACS plots show Uda-SSIEFARL-tetramer^+ve^ cells in the indicated groups of zebrafishes. D. Bar diagrams show the frequencies of virus-specific T cells in HSV1 infected zebrafishes at different time points. E. The specificity of antiviral response in zebrafish was measured *in vitro* by pulsing the CFSE labelled splenocytes from control and the HSV1 infected zebrafishes at 4dpi with SSIEFARL peptide. Representative FACS plots show the frequencies of proliferating splenocytes in response to the peptide stimulation. F. Bar diagram show the cumulative data for the frequencies of proliferating cells. To determine the level of significance between samples obtained from different groups, Student t test and ANOVA were used. The results shown represent mean ± SEM. The level of significance are represented as *, p<0.05, **, p<0.01 and ***<0.001.

Taken together, we demonstrate an expansion of HSV1-specific CD8^+^ T cells in the infected mice. The expanded cells undergo contraction and generate a memory pool that can be recalled by a secondary infection of zebrafish with HSV1 as observed in higher vertebrates.

### 4. Assessing the functionality and migration of HSV1 specific CD8^+^ T cells in zebrafish

We assessed the functionality of HSV1 specific CD8^+^ T cells by measuring their expression of effector molecules, cytolytic activity and the migratory potential. We measured mRNA for the expression of signature molecules such as CD3ε, CD8α and the effector cytokines in the splenocytes, FACS-sorted lymphocytes as well as the Uda-SSIEFARL-tetramer^+ve^ cells from spleens of HSV1 infected zebrafishes (Fig 5A). As compared to the normalised mRNA expression in splenocytes, lymphocytes expressed ∼20 and 40-fold higher levels of CD3ε and CD8α, respectively which indicated that the gated lymphocytes were enriched in CD8^+^ T cells (Fig 5B). The sorted lymphocytes (Fig 5A-B) and the tetramer^+ve^ cells (Fig 5C-D) from the infected fishes expressed upto ∼2 and 5 fold increased levels of IFNγ1 at 2 and 4dpi, respectively (Fig 5A-C). Thereafter, the expression of IFNγ1 decreased to reach at basal level by 7dpi (Fig 5C). A similar kinetic for the expression of TNF-α was observed but the overall expression levels were much low (Fig 5A-C). The mRNA of IL-15, however, decreased at 2dpi but reached to the levels of the uninfected animals at 4dpi (Fig 5A-C). The expression of CD3ε, IFNγ1, TNFα and IL-15 were measured in the Uda-SSIEFARL-tetramer^+ve^ cells (Fig 5D and E). The gated lymphocytes were FACS-sorted into positively and negatively stained populations by Uda-SSIEFARL-tetramer staining and analysed for measuring the expression of CD3ε, IFNγ1, TNFα and IL-15. The tetramer positive cells as compared to the negatively stained cells expressed ∼30, 3, 2.5-fold higher expression levels of CD3ε, IFNγ1, TNFα but 1.5-fold reduced levels of IL-15 (Fig 5D and E). Therefore, the HSV1 reactive T cells of zebrafish upregulated effector molecules following infection.

**Figure 5.**
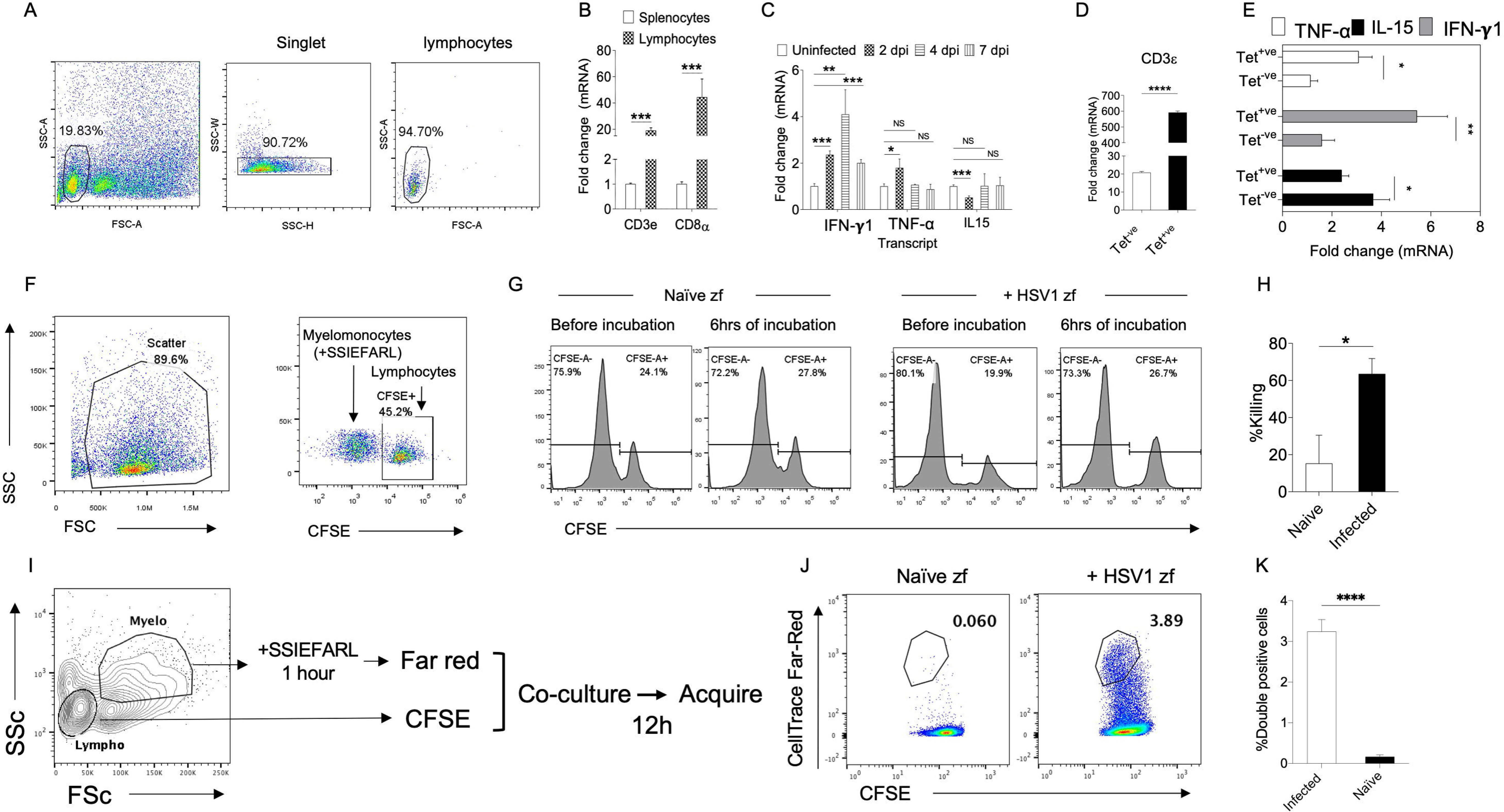
Determining the functionality of HSV1 specific CD8^+^ T cells expanded in the infected zebrafishes. A-E. Gated lymphocytes or the Uda-SSIEFARL-tetramer^+ve^ cells were FACS-sorted from HSV1 infected zebrafishes at 5dpi and the mRNA expression of the indicated genes was measured by quantitative real time PCR. A. Sorting strategy is shown by representative FACS plots. B. The expression of CD3ε and CD8α in the splenocytes or the sorted lymphocytes is shown by bar diagrams. C. Bar diagrams show the expression of IFN-γ1, TNF-α and IL-15 at different time points in the FACS sorted lymphocytes. D and E. FACS-sorted Uda-tetramer positive and tetramer negative cells were analyzed for measuring the mRNA expression of CD3ε, IFN-γ1, TNF-α and IL-15 at 5dpi. Bar diagrams show the expression levels of the indicated genes. To determine the level of significance between samples student t test and ANOVA were used. The results shown are Mean ± SEM values and the level of significance are represented as *, p<0.05, **, p<0.01 and ***<0.001. F-H. *In vitro* cytolytic activity of the HSV1 expanded lymphocytes was assessed by co-culturing the SSIEFARL-peptide pulsed FACS-sorted myelomonocytes and the CFSE labelled lymphocytes from the HSV1 infected and uninfected controls. F. Representative FCAS plots show the gating for the sorted cells (left panel) and the admixed population of the SSIEFARL-peptide pulsed FACS-sorted myelomonocytes as targets and the CFSE labelled lymphocytes (right panel) as the effectors for their identification in the co-cultures. G. Representative histograms show the relative abundance of targets and effectors for calculating killing activity in uninfected controls and the HSV1 infected animals. H. Percentage killing of sorted lymphocytes from naïve and the HSV1 infected zebrafishes is shown by bar diagrams. I-K. The interaction between HSV1 infection expanded lymphocytes with the SSIEFARL-peptide displaying cells was analyzed. I. FACS-sorting strategy of myelomonocytes and lymphocytes is shown. The peptide pulsed myelomonocytes (targets) were labelled with CellTrace Far Red and the lymphocytes were labelled with CFSE. The two cell types were co-cultured for 12 hours and subsequently analyzed for the formation of conjugates. J. Representative FACS plots showing the gated CFSE labelled lymphocytes isolated from uninfected and the HS1 infected zebrafishes are shown. K. The frequencies of conjugates of double positive cells for CellTrace Far Red and CFSE are shown by bar diagram. The results shown by Mean ± SEM. The levels of statistical significance were determined by Student t test and are represented as *, p<0.05, **, p<0.01 and ***<0.001.

One of the critical functions associated with effector CD8^+^ T cells is their ability to kill target cells that can be assessed by co-culturing peptide pulsed targets and lymphocytes from HSV1 infected animals. Based on their scattering characteristics, we FACS purified myelomonocytes and lymphocytes from uninfected controls and the HSV1 infected zebrafishes (Fig 5F). The myelomonocytes were pulsed with SSIEFARL-peptide to serve as the targets. Lymphocytes were labelled with CFSE for their identification in the co-cultures with the targets (Fig 5F). The prepared targets and the effectors from the same animal were admixed in 5:1 ratio and co-incubated *ex vivo* for 6 hrs. Thereafter, the cytolysis by lymphocytes was measured. Lymphocyte population from the infected zebrafishes (65 ± 7%) showed ∼5-fold higher cytolytic activity as compared to those from the uninfected (14 ± 6) zebrafishes in the cocultures (Fig 5G and H).

APCs displaying cognate peptide via their MHC class I molecules interact with the specific TCRs of CD8^+^ T cells leading to the formation of transient conjugates. Such conjugates not only induce activation of T cells but also required for T cell mediated cytolysis. Therefore, a quantification of such conjugates can be used as a surrogate for measuring antigen-specific interactions. A scheme for the experimental procedure is shown in Fig 5I. From naïve and HSV1 infected zebrafish, the FACS-sorted myelomonocytes were pulsed with SSIEFARL peptide and labelled with CellTrace Far red dye while the lymphocytes were labelled with CFSE. Both the cell types isolated from the same zebrafish were admixed and incubated for 12hrs to assess the formation of conjugates by quantifying dual positive cells for CellTrace Far red and CFSE. The gated lymphocytes generated such conjugates more frequently (60-fold) from the HSV1 infected zebrafishes (3.6%) as compared to those obtained from the uninfected zebrafishes (0.06%) suggesting that the HSV1 infected zebrafishes expanded gB-SSIEFARL reactive CD8^+^ T cells which engaged with the peptide pulsed myelomonocytes (Figure 5J and K).

Zebrafish as a model organism offers *in vivo* visualization of cellular dynamics. We, therefore, measured whether or not virus-specific CD8^+^ T cells could be visualised at infected tissue upon their adoptive transfer using confocal microscopy and *in vivo* imaging. Zebrafishes subcutaneously infected with HSV1 were transferred with CFSE labelled Uda-SSIEFARL-tetramer^+ve^ T cells via intracardiac route. The recipient animals were observed by *in vivo* imaging and fluorescent microscopy. *In vivo* imaging revealed that the transferred cells exit blood vessels, alter their trajectory to recruit to the infection sites (Fig 6A and B). The confocal images of tissue sections showed that several CFSE labelled Uda-SSIEFARL-tetramer^+ve^ cells were juxtaposed to the infected cells as the staining of tetramers was polarised towards the contact points with the adjacent cells (Fig 6C). These results indicated the migration of virus-specific lymphocytes to tissue sites and their potential role in the viral clearance by cytolysis. Taken together our results showed that zebrafish as a model for investigating the dynamics of antigen-specific CD8^+^ T cells responses can be explored further.

**Figure 6.**
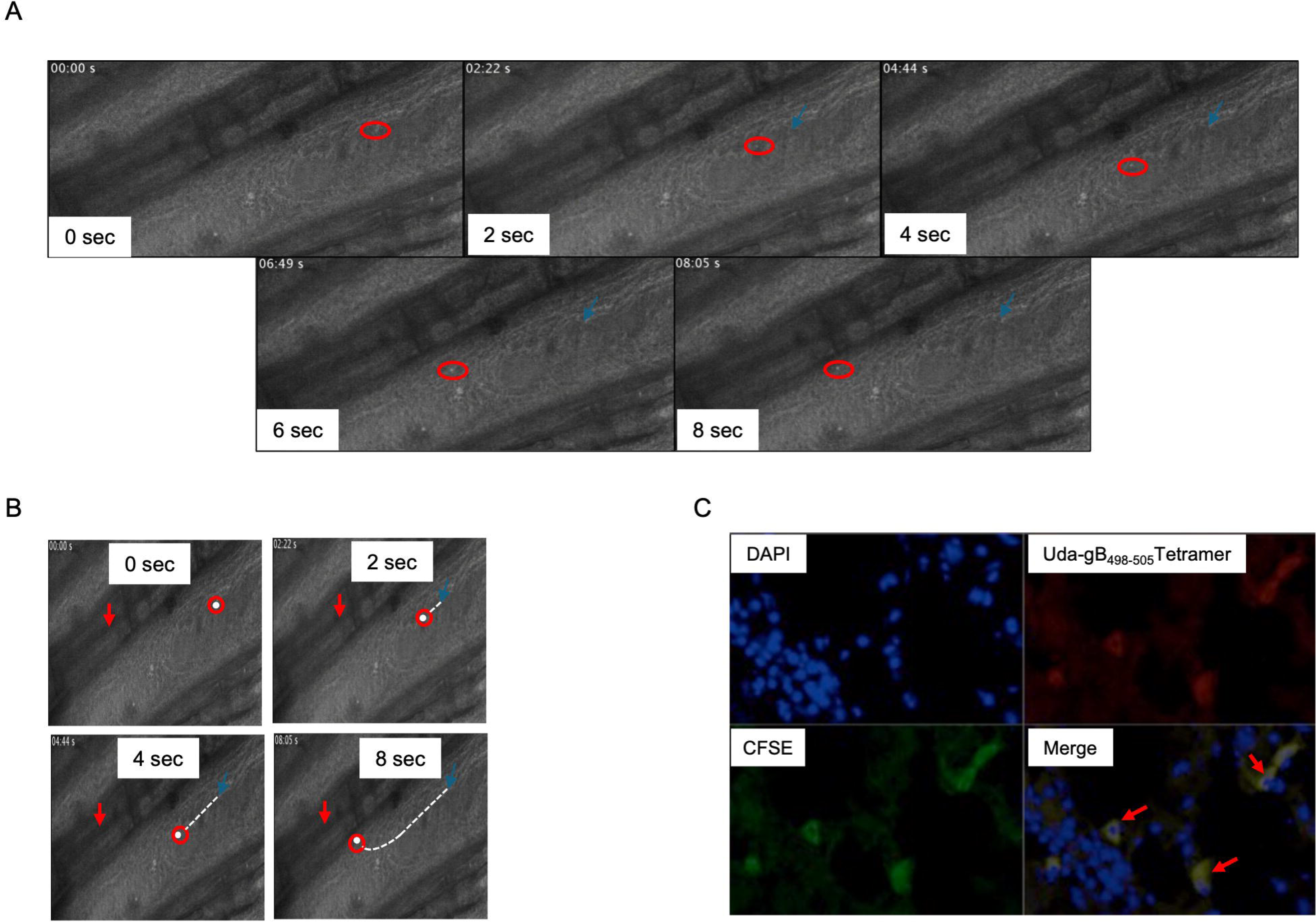
Migratory potential of the tetramer^+ve^ was observed by adoptive transfer of Uda-SSIEFARL-tetramer^+ve^ cells in subcutaneously infected zebrafishes. A-B. Snap shots of video showing the movement of transferred CFSE labeled Uda-SSIEFARL-tetramer^+ve^ cells towards the site of HSV1 infection (marked by a downward red arrow) is shown. The white dot represents the Uda-SSIEFARL-tetramer^+ve^ cell and the dotted lines represent the path of the cell tracked throughout the video. The starting point is represented by a blue arrow. C. Confocal images show the presence of Uda-SSIEFARL-tetramer^+ve^ cells at the site of HSV1 infection. The localization of virus-specific cells at the HSV1 infected sites is shown. The focal and polarized staining of tetramer is shown in proximity of the infected cells.

## Conclusions

The primary objective of this study was to identify and characterize virus-specific T lymphocytes in zebrafishes experimentally infected with HSV1. We demonstrated HSV1 infectivity of adult zebrafishes and the establishment of latency in neuronal tissues by measuring the expression of latency associated transcripts. Such experiments will be useful in further investigating cellular and molecular events in the establishment, maintenance of the viral latency and factors responsible for the reactivation of virus. Furthermore, how antigen-specific T lymphocytes contribute to shaping the balance between lytic and a latent mode of life cycle by the virus could be analysed in a more tractable animal model. We also measured the responsiveness of immune cells to HSV1 antigens and characterised virus-specific cytotoxic T cells by using the generated class I MHC (Uda) tetramers. Furthermore, *in vivo* imaging of virus-specific CD8^+^ T cells and their tissue localization was also assessed in HSV1 infected zebrafishes. With the availability of Uda-SSIEFARL-tetramers, optimal potential of zebrafish as a model can be harnessed for studying cellular dynamics during virus and host interaction. That an immunodominant peptide of HSV1 restricted by H-2K^b^ in C57BL/6 mice could be displayed by Uda class I MHC molecule of zebrafish might suggest for the promiscuity of peptide binding or an existing conserved antigen processing and presentation pathways existing in the two species that diverged ∼445 million years ago in their evolutionary history.

## Materials and methods

### Zebrafish husbandry and their inoculation with HSV1

Zebrafishes were obtained from local vendors and were bred in-house in clean rooms of the animal facility for multiple generations at 26 ± 2°C. The animals bred for 8 generations were used for experimentation. HSV1 was cultivated and titrated using Vero cells as described earlier^40^. Zebrafishes were infected intraperitoneally by injecting 2×10^6^ pfu of HSV1 in 10μl volumes. Spleen, liver and brain tissues were isolated at different days post infection (dpi) and processed to prepare single cell suspensions for further analysis.

### Cloning, expression and purification of zebrafish class I MHC molecule and β2 microglobulin

Total RNA from zebrafish liver was isolated using trizol method and cDNA was synthesised using oligo d(T) and random hexamer primers. The Uda genes of heavy chain of class I MHC molecule and β2 microglobulin were PCR amplified. The sequences of forward and reverse primers used were as follows: For amplifying Uda heavy chain, the primers used were, FP; 5’GGAATTCCATATGGGTACACACTCTCTGAGATACTTCTACACTGCTG3’ and RP 5’ATGGGATCCTA**<u>ATCGCGCAGCTCCATCTTCATAGCCTCGAAGATACCACCCAG</u>**GGGAACAGAAACTTCGTTTGTCC3’. The bold underlined nucleotide sequence encodes for the biotin acceptor peptide (BAP). The primers used for amplifying β2 microglobulin were, FP: 5’GGAATTCCATATGAAAGTCTCCACTCCGAAAGTTCATGTGTACAG3’ and RP: 5’ACGCGGATCCTTACATGTTGGGCTCCCA3’. The PCR product of both the Uda heavy chain and β2 microglobulin were generated as truncated products with the deleted the signal sequences. The cytoplasmic and transmembrane domains for Uda heavy chain were also removed for generating the monomer. In addition to cloning and expressing the truncated product, full length sequence for Uda heavy chain was amplified and cloned separately. The PCR products were cloned in pET22b vector. The plasmid encoding for Uda heavy chain and β2 microglobulin were transformed in BL21 strain of *E. coli* for proteins expression. The protein products were obtained as inclusion bodies.

### Generation of Uda-SSIEFARL-tetramers for zebrafish

The Uda monomers were generated using a rapid dilution method as per the protocol described elsewhere^33^. The composition of refolding buffer was 100mM Tris, 400mM L-Arginine-HCl, 2mM EDTA, 0.5mM oxidized glutathione, 5mM reduced glutathione and 1mM phenylmethyl sulphonyl fluoride (PMSF). Briefly, the peptide, β2 microglobulin and Uda heavy chain were injected sequentially in the rapidly stirred refolding buffer. Different peptides such as SSIEFARL of HSV1 gB, a UV photocleavable peptide, FAPG(Anp)YPAL of Sendi virus and SIINFEKL of Ova protein were used for evaluating the refolding and the generation of Uda monomers. The peptide, β2 microglobulin and the Uda heavy chain were injected following the same sequence for three times with an interval of 12 hours. After 48 hrs of refolding, the refolding reaction mix was filtered and concentrated using 10kDa cutoff Centriprep^®^ filters in a final volume of 1 ml. The concentrated solution was then subjected to size exclusion chromatography using Sephadex S200 columns and fractions of 0.5ml were collected using AKTA pure^TM^. The desired fractions of class I MHC monomers were collected and concentrated. The biotinylation using BirA was achieved by incubating the monomers overnight at room temperature. After biotinylation, monomers were again subjected to size exclusion chromatography. The purified biotinylated class I MHC monomers were collected in different fraction, pooled, concentrated and stored at −80°C until further use. The biotinylation was confirmed by western blotting using conjugate of streptavidin and horse radish peroxidase (streptavidin-HRP). For tetramerizing the monomers, 1nM of streptavidin-phycoerythrin (PE) was added in 10 different aliquots, each separated by 30 minutes, with 4nM of Uda-SSIEFARL monomers at 4°C in dark.

### Flow cytometric analysis of zebrafish immune cells

Spleens, liver and kidneys were collected from control and HSV1 infected zebrafishes at different dpi. Single cell suspensions were prepared for all the extracted organs. RBCs lysis was performed using RBC lysis buffer. The prepared cells were surface stained and analysed using flow cytometry. For quantifying antigen-specific T cells, the cell suspensions from different lymphoid tissues were incubated with PE labelled Uda-SSIEFARL-tetramer at 4°C for 30 min and after three washes, the tetramer stained cells were acquired and analysed by flow cytometery.

### *In vitro* proliferation of virus-specific CD8^+^ T cells

Splenocytes were collected from naïve and HSV1 infected zebrafish at different dpi. The cells from individual zebrafish were labelled with 1µM CFSE and incubated for different durations in the presence or absence of one multiplicity of infection of the UV-inactivated HSV1 virus or different concentrations of SSIEFARL peptide (0.01 to 1μM). The extent of cell proliferation was measured by quantifying the cells that diluted their CFSE content at different time points using flow cytometry.

### Quantitative PCR for effector molecules and viral load determination

Total gated lymphocytes or the Uda-SSIEFARL-tetramer^+ve^ cells were FACS sorted at different time points post HSV1 infection. The total RNA was isolated form the sorted cells using trizol method following the manufacturer’s protocol. DNase treatment was given to remove any DNA contamination. 25ng of total RNA was converted into cDNA using SuperScript IV cDNA synthesis kit from Invitrogen (Cat. No. 18091050). Quantitative PCR (qPCR) was performed using SYBR Green qPCR kit (Thermofisher, Cat. F416L) and a QuantStudio Real-Time PCR system from Thermofisher. The primers used for amplifying zebrafish genes and the product size were as follows: β-Actin (FP: 5’ CGA GCA GGA GAT GGG AAC C 3’ & RP: 5’ CAA CGG AAA CGC TCA TTG C 3’) = 102 bp, CD3ε (FP: 5’ ACA GCG TTT CCA TCC TTT CG 3’ & RP: 5’ GCT CAT CTC CAT CAC CCA CA 3’) = 116 bp, CD8α (FP: 5’ GGA GCA AAG CCC ATC TTG 3’ & RP: 5’ GTG GGG ACA TCG TCT TGT 3’) =132 bp, IFN-γ1 (FP: 5’ TAT GGG CGA TCA AGG AAA AC 3’ & RP: 5’ GCC GTC TCT TGC GTT CTT TA 3’) = 120 bp, TNF-α (FP: 5’ AGG CTG CCA TCC ATT TAA CA 3’ & RP: 5’ CAA GCC ACC TGA AGA AAA GG 3’) = 95 bp, IL-15 (FP: 5’ TGT GCT TTG AGA ATC ACA TGG 3’ & RP: 5’ TGT TCT GGA TGT CCT GCT TG 3’) = 121 bp, HSV1 DNA polymerase (UL30) catalytic subunit (gene id 2703462) (FP: 5’ CAT CAC CGA CCC GGA GAG GGA C 3’ & RP: 5’ GGG CCA GGC GCT TGT TGG TGT A 3’ and major LAT (FP: 5’ CCA GGC AGT AAG ACC CAA GC 3’ & RP: 5’ GGC CGG TGT CGC TGT AAC 3’). The expression levels of mRNA for different molecules were calculated by using 2^−ΔΔ^ CT values. The reaction conditions used for qPCR were: initial denaturation (95°C for 7min), denaturation (95°C for 10sec) then annealing and extension (60°C for 30sec) for total 40 cycles, followed by a melt curve analysis. For viral load determination, total RNA was isolated from spleens of uninfected and zebrafishes infected with HSV1 at different time intervals i.e. 0, 24, 48 and 72 hrs post infection (hpi).

### Viral load determination

HSV1 infected and control zebrafishes were euthanised, dipped in 70% ethanol and extensively washed with sterile PBS. Spleen as well as the rest of the body of the animals were collected and homogenised in 1 and 2 ml serum free cold DMEM, respectively using a tissue rupture at 4°C. The homogenates were centrifuged at 5000 rpm at 4°C for 15min and the supernatants were collected to prepare serial dilutions in serum free DMEM. A volume of 300µl of the diluted supernatants were added to the monolayers of Vero cells. The viral loads as plaque forming units in the organs were calculated as per the following formula:

Pfu/ml = Number of pfu/the volume added (0.3 ml) x dilution of supernatant used
Pfu/spleen = pfu/ml x1
Pfu/whole fish = (pfu/ml) x 2

### *In vitro* CTL assays

Lymphocytes and myelomonocytes were sort purified from zebrafish spleens at 4 dpi. Lymphocytes were stained with CFSE and the myelomonocytes were pulsed with 10ng/ml of SSIEFARL peptide. Lymphocytes and myelomonocytes (target cells) were then mixed in a 1:5 ratio. The frequencies of CFSE^+ve^ and CFSE^−ve^ was measured immediately after the addition of two cell populations or after 6hrs of co-incubation. Killing activity was calculated as the following:-

Target to Lymphocyte ratio (A) = % Target cells at 6h (CFSE^−ve^) / % Lymphocytes at 6h (CFSE^+ve^).
Ratio (B)= A_infected zf_ /Average A _uninfected zf_
% killing = [1-B] × 100
A negative value of % killing was considered as zero, i.e., no killing.

### Analysing conjugates of lymphocytes and the peptide pulsed targets

The FACS-sorted myelomonocytes from naïve and HSV1 infected zebrafish were pulsed with SSIEFARL peptide (10ng/ml) and labelled with CellTrace Far red dye while the lymphocytes were CFSE labelled. Both the cell types isolated from the same zebrafish were admixed and incubated for 12hrs to allow for the formation of conjugates. The cells positively stained for CellTrace Far red and CFSE were quantified by flow cytometry.

### Confocal microscopy

Zebrafishes were infected then sacrificed at 4dpi. Spleen samples were collected and single cell suspension was made. FACS sorted Uda-SSIEFARL-tetramer positive cells labelled with CFSE were transferred into naïve animals through intracardial or retro-orbital plexes routes. Zebrafishes were injected HSV1 in the dorsal side by intramuscular injections and the fishes were euthanised after 30 minutes of the injections. The animals were then fixed in 4% parafromaldehyde prepared in 10mM PBS at 4°C. The samples were then dehydrated in a sucrose gradient from 5 to 20% in 10mM PBS at room temperature and embedded in OCT compound in tissue blocks. The blocks were frozen at −80°C. Six-micrometer tissue sections were cut using a Leica cryotome, and the slides were stored at 20°C until further use. The sections were then stained overnight at 4°C with Uda-gB_498-505_-tetramer-PE. After two washes with PBS, the slides were stained with 4′,6-diamidino-2-phenylindole (DAPI) dye. The stained sections were dried at room temperature and coverslips were mounted using a mounting medium. Images were acquired using Nikon A1 confocal microscope and analysed by ImageJ software.

### Statistical analysis

To determine the level of significance between samples obtained from different groups, student t test and ANOVA were used. The results shown represent mean ± SD. The level of significance are represented as *, p<0.05, **, p<0.01 and ***<0.001.

## References

1. Ma D, Zhang J, Lin HF, Italiano J, and Handin RI (2011). The identification and characterization of zebrafish hematopoietic stem cells. Blood, The Journal of the American Society of Hematology, 118(2), 289–297.

2. Stachura DL, Reyes JR, Bartunek P, Paw BH, Zon LI, and Traver D (2009). Zebrafish kidney stromal cell lines support multilineage hematopoiesis. Blood, The Journal of the American Society of Hematology, 114(2), 279–289.

3. Stachura DL and Traver D (2016) Cellular dissection of zebrafish haematopoiesis. in Methods in cell biology 133, 11–53.

4. Danilova N, and Steiner LA (2002). B cells develop in the zebrafish pancreas. Proc. of the Natl. Acad of Sci., 99(21), 13711–13716.

5. Kari G, Rodeck U and Dicker AP (2007) Zebrafish: An Emerging Model System for Human Disease and Drug Discovery. Clin. Pharmacol. Ther. 82, 70–80.

6. Kaur J, Sohal IS, Singh H, Gupta NK, Sehrawat S, Puri S, Bello D and Khatri M (2022) Toxicity screening and ranking of diverse engineered nanomaterials using established hierarchical testing approaches with a complementary in vivo zebrafish model. Environmental Science: Nano. 9(8), 2726–2749.

7. Malik JA, Nanda S, Zafar MA, Sehrawat S and Agrewala JN (2023) Influence of chronic administration of morphine and its withdrawal on the behaviour of zebrafish. J. Biosci. 48, 33

8. Kaur M, Dubey A, Khatri M and Sehrawat S (2019) Secretory PLA2 specific single domain antibody neutralizes Russell viper venom induced cellular and organismal toxicity. Toxicon. 172,15–18.

9. Lieschke GJ and Currie PD (2007) Animal models of human disease: Zebrafish swim into view. Nat. Rev. Genet. 8, 353–367.

10. Trede NS, Langenau DM, Traver D, Look AT and Zon LI (2004) The use of zebrafish to understand immunity. Immunity. 20, 367–79.

11. Meeker ND and Trede NS (2008). Immunology and zebrafish: Spawning new models of human disease. Dev. Comp. Immunol. 32, 745–757.

12. Yoder JA, Nielsen ME, Amemiya CT and Litman GW (2004) Zebrafish as an immunological model system. Microbes Infect. 4, 1469–78.

13. Trede NS, Langenau DM, Traver D, Look AT and Zon LI (2004) The use of zebrafish to understand immunity. Immunity 20, 367–79.

14. Renshaw SA and Trede NS (2012) A model 450 million years in the making: zebrafish and vertebrate immunity. Dis. Model. Mech. 5, 38–47.

15. Lugo-Villarino G, Balla KM, Stachura DL, Banuelos K, Werneck MBF and Traver D (2010) Identification of dendritic antigen presenting cells in the zebrafish. Proc. Natl. Acad. Sci. U. S. A 107, 15850–5.

16. Kasheta M, Painter CA, Moore FE, Lobbardi R, Bryll A, Freiman E, Stachura D, Rogers AB, Houvras R, Langenau DM and Ceol CJ (2017) Identification and characterisation of T reg-like cells in zebrafish. J. Exp. Med. 214, 3519–3530.

17. Wan F, Hu CB, Ma JX, Gao K, Xiang LX and Shao JZ (2017) Characterisation of γδ T cells from zebrafish provides insights into their important role in adaptive humoral immunity. Frontiers in immunology, 7, 675, 1–18.

18. Neely MN, Pfeifer JD and Caparon M (2002). Streptococcus-zebrafish model of bacterial pathogenesis. Infection and immunity, 70(7), 3904–3914.

19. Novoa B and Figueras A (2012). Zebrafish: model for the study of inflammation and the innate immune response to infectious diseases. In Current topics in innate immunity II (pp. 253–275). Springer, New York, NY.

20. Gomes MC and Mostowy S (2019) The case for modelling human infection in zebrafish. Trends in microbiology. 28(1), 10–18.

21. Neely MN (2017). The zebrafish as a model for human bacterial infections. In Bacterial Pathogenesis (pp. 245–266). Humana Press, New York, NY.

22. Rakus K, Adamek M, Mojżesz M, Podlasz P, Chmielewska-Krzesińska M, Naumowicz K, Kasica-Jarosz N, Klak K, Rakers S, Way K, Steinhagen D and Chadzinska M (2019). Evaluation of zebrafish (Danio rerio) as an animal model for the viral infections of fish. J. of fish dis. 42(6), 923–934.

23. Crim MJ and Riley LK (2012). Viral diseases in zebrafish: what is known and unknown. ILAR journal, 53(2), 135–143.

24. Brown JC (2017). Herpes simplex virus latency: the DNA repair-centered pathway. Adv in virol. 7028194.

25. Sehrawat S, Kumar D and Rouse BT (2019) Herpesviruses: Harmonious pathogens but relevant cofactors in other diseases. Front. in Cell. Infect. Microbiol., 8, 177.

26. Weller SK and Coen DM (2012) Herpes simplex viruses: mechanisms of DNA replication. Cold Spring Harbor perspectives in biology, 4(9), a013011.

27. Nicoll MP, Proença JT and Efstathiou S (2012). The molecular basis of herpes simplex virus latency. FEMS microbiol. reviews, 36(3), 684–705.

28. Burgos JS, Ripoll-Gomez J, Alfaro JM, Sastre I and Valdivieso F (2008) Zebrafish as a New Model for Herpes Simplex Virus Type 1 Infection. Zebrafish. 5, 323–333.

29. Antoine TE, Jones KS, Dale RM, Shukla D and Tiwari V (2014) Zebrafish: Modelling Herpes Simplex Virus Infections. Zebrafish. 11, 17–25.

30. Yakoub AM, Rawal N, Maus E, Baldwin J, Shukla S and Tiwari V (2014) Comprehensive analysis of Herpes Simplex Virus 1 (HSV-1) Entry Mediated by Zebrafish 3-O-Sulfotransferase Isoforms: Implications for the Development of a Zebrafish Model of HSV-1 Infection. J. Virol. 88, 12915–12922.

31. Burnham LA, Jaishankar D, Thompson JM, Jones KS, Shukla S and Tiwari V (2016) Liposome-mediated herpes simplex virus uptake is glycoprotein-D receptor-independent but requires heparan sulfate. Front. Microbiol. 7, 973.

32. Wallace ME, Keating R and Heath WR (1999) The Cytotoxic T-Cell Response to Herpes Simplex Virus Type 1 Infection of C57BL / 6 Mice is Almost Entirely Directed Against a Single Immunodominant Determinant. J. Virol. 73, 7619–7626.

33. Treat BR, Bidula SM, Ramachandran S, St Leger AJ, Hendricks RL and Kinchington PR (2017). Influence of an immunodominant herpes simplex virus type 1 CD8+ T cell epitope on the target hierarchy and function of subdominant CD8+ T cells. PLoS pathog. 13(12), e1006732.

34. McConnell SC, Hernandez KM, Wcisel DJ, Kettleborough RN, Stemple DL, Yoder JA, Andrade J, de Jong JLO (2016) Alternative haplotypes of antigen processing genes in zebrafish diverged early in vertebrate evolution. Proc. Natl. Acad. Sci USA 4;113(34):E5014–E5023

35. Schatz PJ (1993) Use of Peptide Libraries to Map the Substrate Specificity of a Peptide-Modifying Enzyme: A 13 Residue Consensus Peptide Specifies Biotinylation in *Escherichia coli*. Nat. Biotechnol. 11, 1138–1143.

36. John AD, Moss PAH, Goulder PJR, Barouch DH, McHeyzer-Williams MG, Bell JI, McMichael AJ and Davis MM (1996) Phenotypic analysis of antigen-specific T lymphocytes. Science 274(5284), 94–96.

37. Sarkar R, Sharma Y, Jain A, Tehseen A, Singh S and Sehrawat S (2021) A combinatorial in-silico, in-vitro and in-vivo approach to quantitatively study peptide induced MHC stability. Bio-protocol 11 (24), e4255–e4255

